# Role of left motor cortex in post-stroke fatigue: a corticospinal excitability study

**DOI:** 10.1101/2020.12.09.417766

**Authors:** William De Doncker, Annapoorna Kuppuswamy

**Author notes:** Corresponding author: Mr William De Doncker, Box 146, 33 Queen Square, London WC1N 3BG.

## Abstract

**Background:** The neural mechanisms that underlie post-stroke fatigue are poorly understood. Previous work show an inverse relationship between motor cortex excitability and post-stroke fatigue, however, it is unclear if the side of lesion influences this relationship. The left hemisphere plays a dominant role in motor control, therefore we hypothesised that left hemisphere strokes are more likely to show a significant inverse relationship between corticospinal excitability and fatigue.

**Methods:** Resting motor threshold (measure of corticospinal excitability) using transcranial magnetic stimulation was measured in the affected hemisphere of 98 stroke survivors. Fatigue was measured using the fatigue severity scale. The effect of fatigue and hemisphere affected on corticospinal excitability was analysed using a multiple linear regression.

**Results:** A multiple linear regression with trait fatigue as the outcome variable (F_(4,93)_=12.04, p < 0.001, adj R^2^ = 0.313) revealed that RMT was not a significant predictor of FSS-7 (β = −0.063, p = 0.706, CI[-0.394, 0.268]), while the interaction between lesioned hemisphere and RMT was a significant predictor of FSS-7 (β = 0.339, p = 0.039, CI[0.018, 0.659]). The additional explanatory variables of HADS_Depression_ and sex were also significant predictors of FSS-7 (β = 903, p < 0.001, CI[0.584, 1.223] and β = 1.127, p = 0.002, CI[0.425, 1.830] respectively).

**Conclusion:** Lower corticospinal excitability of the left hemisphere may indicate altered perception of effort and reduced sensory attenuation. This provides evidence to support the sensory attenuation model of fatigue.

## Introduction

Fatigue that is chronic is one of the most commonly self-reported symptoms in a number of neurological disorders from stroke to multiple sclerosis (MS) and Parkinson’s disease (Krupp, 2006; Barone *et al*., 2009; Naess *et al*., 2012). Incidence of post-stroke fatigue (PSF) has been reported to be as high as 85% and has a significant impact on disability and quality of life (van de Port *et al*., 2007; Cumming *et al*., 2016). PSF has been identified by stroke survivors as one of their top unmet needs and is a top priority of further research (Kirkevold *et al*., 2012; Pollock *et al*., 2014). One of the difficulties in studying PSF is the significant overlap it shares with other affective symptoms such as depression and anxiety.

Despite the high prevalence of PSF, an understanding of the neural mechanisms that underlie it is currently lacking. We recently proposed that increased perceived effort as result of reduced sensory attenuation (i.e. inability to modulate attention away from predictable sensory input) gives rise to PSF (Kuppuswamy, 2017, De Doncker *et al*., 2020*b*). Slower self-selected ballistic movement speeds, perceived limb heaviness, reduced corticospinal excitability, reduced modulation of corticospinal excitability during movement preparation and poor attention all support the proposed model of PSF (Radman *et al*., 2012, Kuppuswamy *et al*., 2015*b, a, c*, De Doncker *et al*., 2020*a*). However, we have thus far not investigated the influence of side of lesion on neurophysiological parameters in relation to fatigue.

In the normal functioning brain, there is an asymmetry between the two hemispheres within and outside primary motor areas characterized by a net inhibitory dominance from the left hemisphere (Ziemann and Hallett, 2001; Corbetta and Shulman, 2002; Mevorach *et al*., 2006; Giovannelli *et al*., 2009). We have recently showed that the interhemispheric inhibition balance (IIB), a measure of interhemispheric network dynamics, is altered in PSF, with a shift from a net left-to-right inhibition dominance to a net right-to-left inhibition dominance (Ondobaka *et al*., 2019). This shift in IIB has also previously been shown to account for clinical depression, with overall lower corticospinal excitability seen in the left hemisphere compared to the right hemisphere (Lefaucheur *et al*., 2008). Non-invasive brain stimulation studies have shown that IIB has an influence of corticospinal excitability of both hemispheres (Schambra *et al*., 2003). Dysfunctional functional connectivity within sensorimotor networks, particularly of the left hemisphere, appear to also play a crucial role in mediating MS fatigue (Dell’Acqua *et al*., 2010; Cogliati Dezza *et al*., 2014; Chalah *et al*., 2015; Vecchio *et al*., 2017). Targeting the connectivity within these regions using non-invasive brain stimulation has shown to reduce the severity of fatigue (Tecchio *et al*., 2014; Porcaro *et al*., 2019).

Given the directionality of IIB in PSF, the overlap between fatigue and depression and the role of the left hemisphere in mediating fatigue in other neurological disorders, we aimed to investigate the influence of fatigue on corticospinal excitability in stroke survivors with left and right hemisphere strokes. We hypothesised that stroke survivors with left hemisphere strokes will have lower corticospinal excitability while stroke survivors with right hemisphere strokes will have higher corticospinal excitability with increasing severity of fatigue.

## Materials and Methods

### Subjects

This is a single session cross-sectional observational study approved by the London Bromley Research Ethics Committee (REC reference number: 16/LO/0714). Stroke survivors were recruited via the Clinical Research Network from the University College NHS Trust Hospital, a departmental Stroke Database and from the community. All stroke survivors were screened prior to the study based on the following criteria: (1) first-time ischaemic or haemorrhagic stroke; (2) stroke occurred at least 3 months prior to the study; (3) no other neurological disorder; (4) not taking anti-depressants or any other centrally acting medication; (5) no clinical diagnosis of depression with depression scores ≤ 11 assessed using the Hospital Anxiety and Depression Scale (HADS); (6) grip strength and manual dexterity of the affected hand (at least 60% of the unaffected hand) assessed using a hand-held dynamometer and the nine hole peg test (NHPT) respectively; (7) no contraindications to transcranial magnetic stimulation (TMS). Ninety-eight stroke survivors took part in the study (Table 1). All stroke survivors provided written informed consent in accordance with the Declaration of Helsinki.

**Table 1.** Demographics of patients that completed the study. Mean (SD) values are displayed for all continuous variables and number of patients are displayed for categorical variables (Hemisphere affected and gender). P-values indicate the significance of spearman rank correlations between FSS-7 and continuous variables and Wilcoxon rank sum tests comparing the FSS-7 score within categorical values.

| DEMOGRAPHICS | N = 98 | STATISTIC | P-VALUE |
| --- | --- | --- | --- |
| **FSS-7** | 3.83 (1.91) |  |  |
| **HEMISPHERE AFFECTED (LEFT \| RIGHT)** | 51 \| 47 | 1379 | 0.200 |
| **SEX (FEMALES \| MALES)** | 32 \| 66 | 661 | **0.003** |
| **AGE (YEARS)** | 60.93 (12.65) | -0.198 | 0.050 |
| **TIME POST-STROKE (YEARS)** | 4.51 (4.74) | -0.097 | 0.348 |
| **GRIP (% OF UNAFFECTED HAND)** | 91.37 (22.93) | -0.204 | **0.044** |
| **NHPT (% OF UNAFFECTED HAND)** | 88.48 (24.63) | -0.123 | 0.229 |
| **HADS<sub>DEPRESSION</sub>** | 4.83 (3.23) | 0.498 | **<0.001** |
| **HADS<sub>ANXIETY</sub>** | 5.26 (3.93) | 0.367 | **<0.001** |

### Questionnaires

Fatigue was quantified using the fatigue severity scale (FSS-7), a seven-item questionnaire asking for ratings of fatigue ranging from one to seven (strongly disagree to strongly agree) over the preceding week from the day of administration (Krupp *et al*., 1989). An average score of one indicates no fatigue while an average score of seven indicates very severe fatigue. Patients also completed the HADS, a 14-item questionnaire with a depression (HADS_Depression_) and anxiety (HADS_Anxiety_) subscale, prior to the stimulation. A score of 0 to 7 for either subscale could be regarded as being in the normal range, with a score of 11 or higher indicating probable presence of the mood disorder (Snaith, 2003).

### Surface electromyogram and transcranial magnetic stimulation

Recordings were carried out on the first dorsal interosseous (FDI) muscle of the affected hand. Following skin preparation using alcohol swabs, electromyogram (EMG) recordings were obtained from the FDI muscle using surface neonatal prewired disposable electrodes (1041PTS Neonatal Electrode, Kendell) in a belly-tendon montage with the ground positioned over the flexor retinaculum of the hand. The signal was amplified with a gain of 1000 (D360, Digitmer, Welwyn Garden City, UK), bandpass filtered (100-1000 Hz), digitized at 5 kHz (Power1401, CED, Cambridge, UK) and recorded with Signal version 6.04 software (CED, Cambridge, UK). EMG recordings enabled the measurement of motor evoked potentials (MEPs).

A standard monophasic TMS device (Magstin 2002, Magstim, Whitland, Wales) connected to a figure-of-eight coil (wing diameter, 70 mm) was used to stimulate the hand area of the primary motor cortex (M1) in the hemisphere affected by the stroke. The coil was held tangentially on the scalp at an angle of 45o to the mid-sagittal plane to induce a posterior-anterior current across the central sulcus. The subjects were instructed to stay relaxed with their eyes open and their legs uncrossed. The motor ‘hotspot’ of the FDI muscle was determined as follows: the vertex (cross-over point between the mid-point between the two tragi and midpoint between nasion and inion) was marked using a dry wipe marker. Four centimetres lateral and 2 cm anterior from the vertex was then marked on the contralateral hemisphere, which is the approximate location of M1. This was used as a rough guide for a starting point for determining the hotspot for the first dorsal interosseous. At 50% maximal stimulator output (MSO) (or higher or lower in some patients) the coil was moved in 1 cm blocks for ∼2 cm anterior, posterior, lateral and medial to the marked region. Three stimuli were delivered at each spot and the location with the highest average motor evoked potential response was taken as the hotspot.

Resting motor threshold (RMT) was defined as the lowest intensity of stimulation (% MSO) required to evoke a peak-to-peak MEP amplitude at the hotspot of at least 50 μV in a minimum of 5 of 10 consecutive trials while subjects were at rest.

### Statistical Analysis

The association between FSS-7 and time post-stroke, age, grip strength, NHPT, HADS_Depression_and HADS_Anxiety_ was assessed using Spearman’s Rank Correlations. Wilcoxon rank sum tests were used to assess the difference in FSS-7 across different groups divided based on lesioned hemisphere and sex.

To explain individual differences in fatigue, a multiple linear regression was used (RStudio Version 1.2.5033) with RMT and the interaction between lesioned hemisphere (left vs right) and RMT as explanatory variables. The interaction term was included to test for a potential confounding effect of lesioned hemisphere. HADS_Depression_ scores and sex significantly improved the model, assessed using the BIC (lower BIC indicates a better fitting model) and were included as independent explanatory variables. Multicollinearity of the explanatory variables was assessed by computing the variance inflation factor (VIF). Assumptions of normality and homoscedasticity of the residuals for each linear regression model were assessed visually using quantile-quantile normal plots and fitted-versus residual-value plots.

## Results

### Demographics

There was a significant association between FSS-7 and grip strength (rho=-0.204, p=0.044; figure 1C), HADS_Anxiety_ (rho=0.367, p<0.001; figure 1E) and HADS_Depression_(rho=0.498, p<0.001; figure 2B) in the 98 stroke survivors. There was no association between FSS-7 and time post-stroke (rho=-0.097, p=0.348; figure 1A), age (rho=-0.198, p =0.050; figure 1B) and NHPT (rho=-0.123, p=0.229; figure 1D). There was a significant difference in median FSS-7 scores between males and females (W = 661, p=0.003; figure 2C) but no significant difference between left and right hemisphere strokes (W=1379, p=0.200; figure 1F) in the cohort of stroke survivors (demographics in table 1).

**Figure 1.**
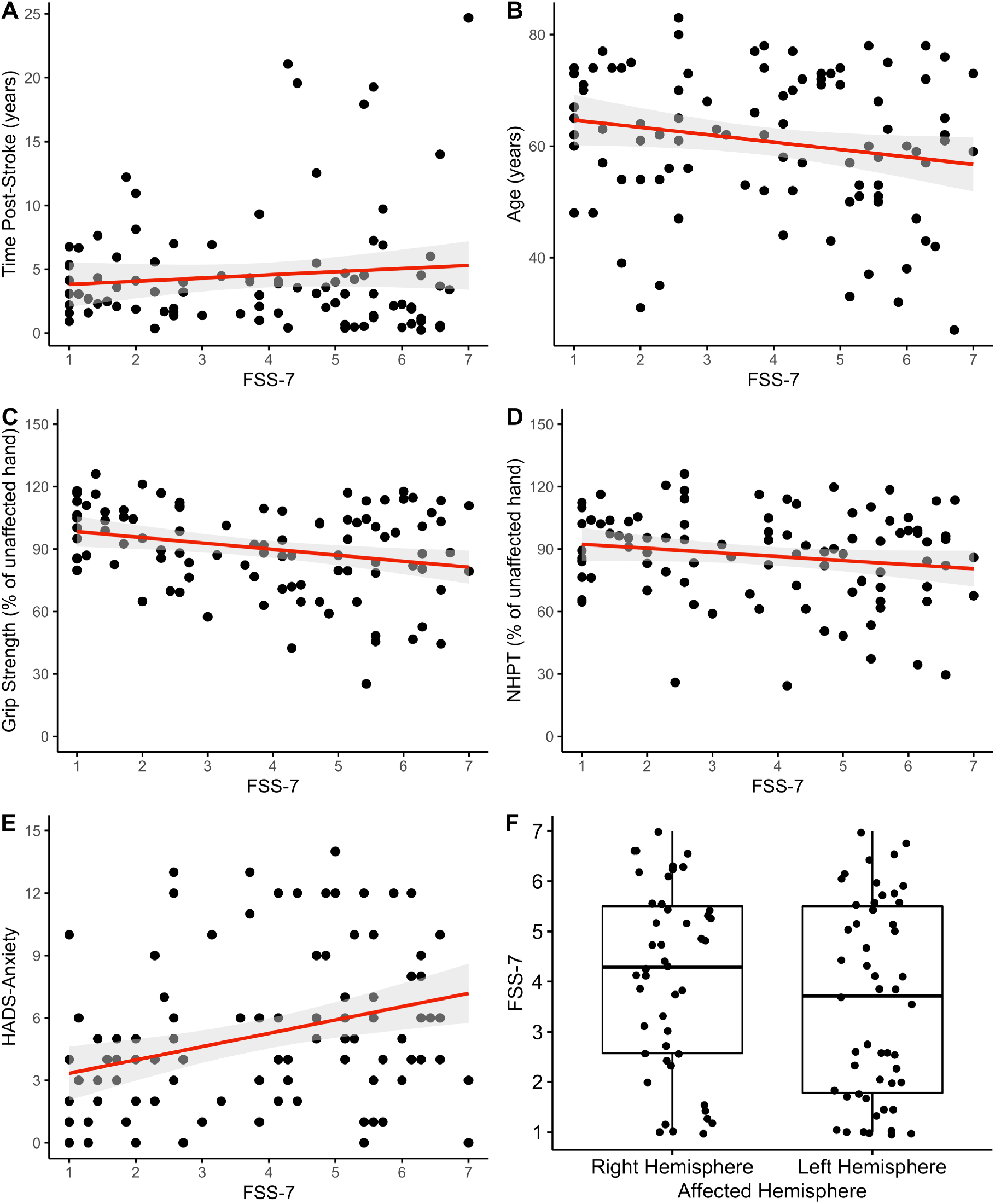
Association between fatigue (FSS-7) and demographic data across the entire cohort of stroke survivors (n = 98). For panel A-E, the red line indicates the regression line and its associated 95% confidence interval in grey for continuous data. For panel F, boxplots displaying the median and interquartile range for categorical data (hemisphere affected).

**Figure 2.**
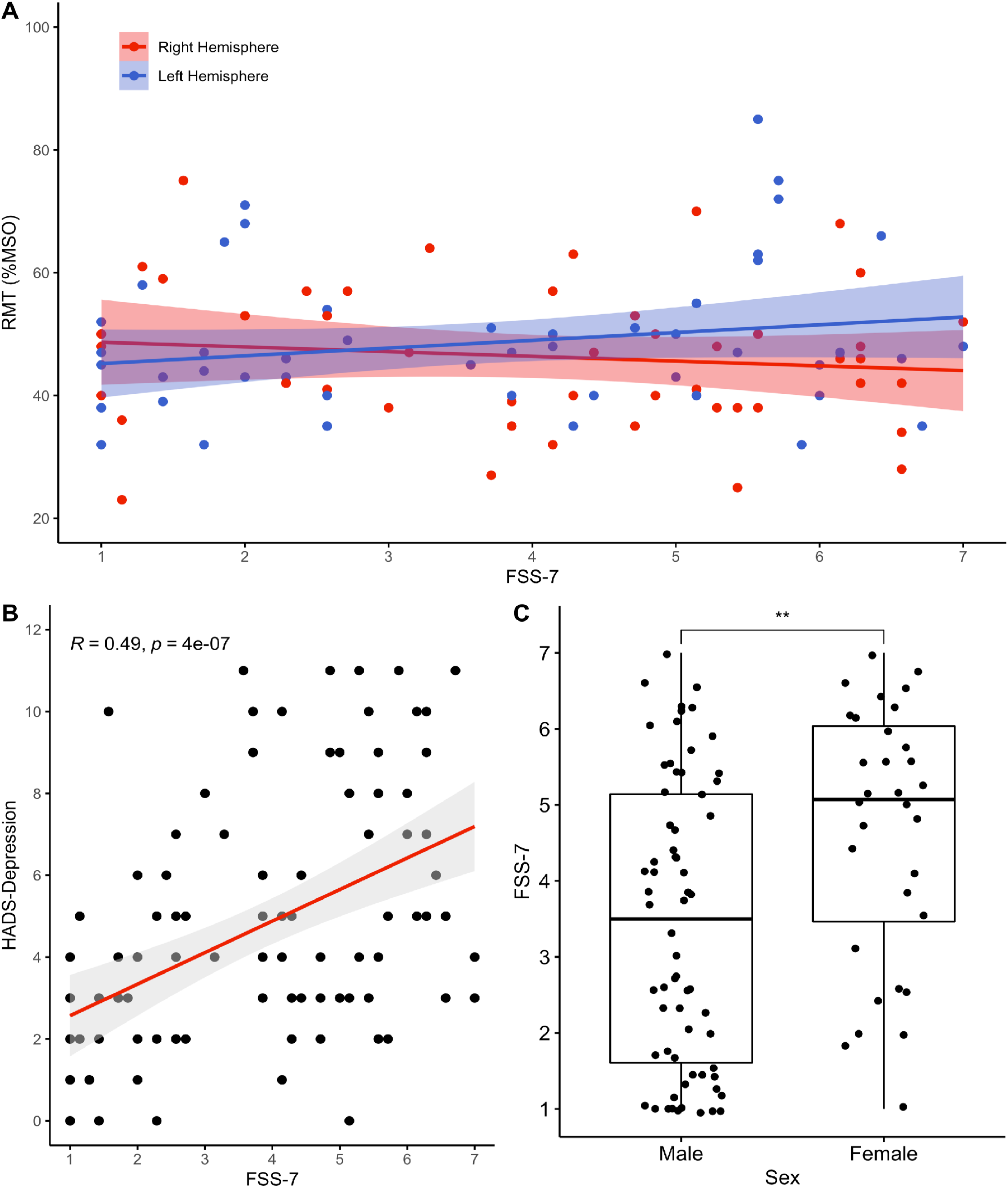
A) The association between resting motor threshold (RMT) and fatigue (FSS-7). Stroke survivors are displayed based on the hemisphere affected by the stroke with right hemisphere strokes in blue and left hemisphere strokes in blue. Regression lines and the associated 95% confidence interval is displayed is shown for both groups. B) Scatter plot with a regression line and 95% confidence interval showing the association between fatigue (FSS-7) and HADS_Depression_. C) Boxplot displaying the effect of sex on fatigue (FSS-7) in the cohort of stroke survivors. ** indicate a significance level of <0.01.

### Resting Motor Threshold

The multiple linear regression equation explained 31.3% of the variance in fatigue (F_(4,93)_=12.04, p<0.001, adj R^2^ = 0.313). RMT was not a significant predictor of FSS-7 (β = −0.063, p = 0.706, CI[-0.394, 0.268]), while the interaction between lesioned hemisphere and RMT was a significant predictor of FSS-7 (β = 0.339, p = 0.039, CI[0.018, 0.659]; figure 2A). The additional explanatory variables of HADS_Depression_(figure 2B) and sex (figure 2C) were also significant predictors of FSS-7 (β = 903, p < 0.001, CI[0.584, 1.223] and β = 1.127, p = 0.002, CI[0.425, 1.830] respectively).

## Discussion

In this study we show that stroke survivors with left hemisphere strokes and high fatigue have lower corticospinal excitability than those with low levels of fatigue and right hemisphere strokes. Stroke survivors with high severity of fatigue scored higher on both the anxiety and depression subscales of HADS and also had less grip strength on their affected hand when compared to their unaffected hand. Fatigue severity was higher in female stroke survivors compared to males.

Corticospinal excitability assessed using TMS resting motor threshold is highly variable in the general population (Wassermann, 2002). The two hemispheres behave differently during motor control. The left hemisphere appears to have a dominant role during motor control characterized by higher corticospinal excitability, broader activating patterns during movement and greater thickness of the motor cortex (Hammond, 2002; Hervé *et al*., 2009; Verstynen and Ivry, 2011; Davidson and Tremblay, 2013; Klein *et al*., 2016). This is in line with the asymmetry in inter-hemispheric connectivity, a net left-to-right inhibitory dominance and a net right-to-left excitatory dominance, seen in normal functioning brains. In PSF, there is a shift of inter-hemispheric effective connectivity within primary motor cortices from a net left-to-right to a net right-to-left inhibitory dominance (Ondobaka *et al*., 2019). In multiple sclerosis fatigue, sensorimotor networks within the left hemisphere show the biggest change following neuromodulation techniques (Porcaro *et al*., 2019). The authors suggest that re-balancing the inter-hemispheric difference between sensorimotor networks in the left and right hemispheres reduces fatigue severity. One would therefore expect to see lower corticospinal excitability in the left hemisphere and higher corticospinal excitability in the right hemisphere of stroke survivors with high severity of fatigue. In line with this prediction, we show stroke survivors with high fatigue and left hemisphere strokes had lower corticospinal excitability than those with low fatigue and right hemisphere strokes.

Corticospinal excitability does not only reflect the output of the motor cortex but also the excitability of inputs that drive the output. The motor cortex has strong anatomical connections with other cortical and subcortical regions such as the pre-motor cortices, supplementary motor areas, cingulate motor areas, basal ganglia and the cerebellum, all of which can modulate corticospinal excitability (Strick *et al*., 1998; Kirimoto *et al*., 2011; Lee *et al*., 2013). Higher motor areas of the left-hemisphere specifically have a dominant role in motor attention and movement selection (Rushworth *et al*., 2003) and disrupting neural activity within the supplementary motor area of the left hemisphere results in a decrease in effort perception (Zénon *et al*., 2015). We recently proposed a model of PSF whereby poor sensory attenuation leads to increased perceived effort and subsequently high fatigue (Kuppuswamy, 2017). In fact, artificially reducing corticospinal excitability by stimulating the primary motor cortex of the left hemisphere using theta-burst stimulation reduces sensory attenuation (Voss *et al*., 2007). Corticospinal excitability is therefore closely associated to motor attention, effort perception and sensory attenuation. The results of the current study, reduced corticospinal excitability of the left hemisphere of stroke survivors with high fatigue, provides evidence to support the sensory attenuation model of fatigue.

The association seen between fatigue and the anxiety and depression subscales of HADS is not surprising. There is significant overlap between affective symptoms in neurological disorders (Cumming *et al*., 2018; De Doncker *et al*., 2018), suggesting that fatigue, anxiety and depression may share common underlying mechanisms resulting in a cluster of symptoms (Ayache and Chalah, 2019). This is also evident from similar findings of lower corticospinal excitability in the left hemisphere of patients with clinical depression (Lefaucheur *et al*., 2008). It is important to note however, that despite the overlap, fatigue can exist independently of other affective symptoms (van der Werf *et al*., 2001). This highlights the importance of using a strictly controlled patient cohort when trying to draw conclusions from studies and attempting to develop a mechanistic understanding of affective symptoms.

The finding of greater fatigue in women than men has previously been reported in the stroke population (Schepers *et al*., 2006; Mead *et al*., 2011). Whether the imbalance is reflective of report bias with men considering less acceptable to report fatigue related symptoms or a difference in physiology remains unknown. It is important to consider that in the current cohort of stroke survivors there were twice as many males than females. The difference in fatigue between males and females observed in this study may therefore be a sampling issue. Our finding that higher levels of upper limb impairment, indicated by lower grip strength of the affected hand compared to the unaffected hand, are associated with PSF also supports previous studies (Choi-Kwon *et al*., 2005). It is important to note that all stroke survivors that took part in the study were fully independent and did not consider themselves to have a physical disability. Despite having lower grip strength, the manual dexterity of those with high fatigue was relatively high as stroke survivors that had less than 60% grip strength and dexterity on the affected hand were excluded from the study. Therefore, we do not consider muscle weakness as a cause of reported fatigue.

Our results take us a step towards developing a mechanistic understanding of post-stroke fatigue. The relationship between corticospinal excitability of the affected hemisphere and self-reported fatigue may be influenced by other variables than previously suggested (Kuppuswamy *et al*., 2015*c*). In this study we show that the hemisphere affected significantly contributes to the association between corticospinal excitability and fatigue. Identifying the functional role of the left hemisphere in mediating fatigue across a range of neurological and psychiatric disorders makes it a potential target for developing effective interventions of fatigue.

## Acknowledgments

We thank Mr Cameron Cook and the Clinical Research Network for their help with recruitment. We thank our lab manager Mr Paul Hammond for the technical support throughout the project. We extend our heartfelt thanks to all our stroke survivor participants in this study without whose enthusiasm and commitment this study would not have been possible.

## Funding

This work was supported by the Wellcome Trust (202346/Z/16/Z).

## Conflict of Interest

The authors report no conflict of interest.

## Data Availability Statement

Data is available upon reasonable request from William De Doncker

## Abbreviations

MS: Multiple sclerosis
PSF: post-stroke fatigue
IIB: interhemispheric inhibition balance
HADS: hospital anxiety and depression scale
NHPT: nine-hole peg test
TMS: transcranial magnetic stimulation
FSS-7: fatigue severity scale
FDI: first dorsal interosseous
EMG: electromyogram
MEP: motor evoked potential
M1: primary motor cortex
MSO: maximum stimulator output
RMT: resting motor threshold.

## Notes

### Competing Interest Statement

The authors have declared no competing interest.

